# OGG1 inhibitor TH5487 alleviates allergic airway inflammation in mice

**DOI:** 10.1101/2022.05.17.492235

**Authors:** Lloyd Tanner, Jesper Bergwik, Ravi KV Bhongir, Lang Pan, Caijuan Dong, Christina Kalderén, Thomas Helleday, Istvan Boldogh, Mikael Adner, Arne Egesten

**Affiliations:** Respiratory Medicine, Allergology, and Palliative Medicine, Department of Clinical Sciences Lund, Lund University and Skåne University Hospital, Lund, Sweden, SE-221 84; Department of Microbiology and Immunology, University of Texas Medical Branch at Galveston, Galveston, TX77555, USA; Unit of Experimental Asthma and Allergy Research, Institute of Environmental Medicine (IMM), Karolinska Institutet, Stockholm, Sweden, SE-171 77; Science for Life Laboratory, Department of Oncology-Pathology, Karolinska Institutet, SE-171 76 Stockholm, Sweden; Oxcia AB, Norrbackagatan 70C, SE-113 34 Stockholm; Weston Park Cancer Centre, Department of Oncology and Metabolism, University of Sheffield, Sheffield S10 2RX, UK

**Keywords:** Allergic asthma, OGG1 inhibitor, macrophage polarization, NF-κB, T_H_2 cytokines

## Abstract

Allergic asthma is a complex disease characterized by dyspnea, coughing, chest tightness and airway remodeling, for which there is no cure and is symptomatically treated with inhaled β2-agonist and/or corticosteroids. Molecular mechanisms underlying its complex pathogenesis are not fully understood. However, the 8-oxoguanine DNA glycosylase-1 (OGG1), a DNA repair protein may play a central role, as OGG1 deficiency decreases both innate and allergic inflammatory responses. In this study, administration of TH5487 to mice with OVA-induced allergic airway inflammation significantly decreased goblet cell hyperplasia and mucus production. TH5487 treatment also decreased levels of activated NF-κB and expression of proinflammatory cytokines resulting in significantly lower recruitment of eosinophils and other immune cells to the lungs. Gene expression profiling of asthma and allergy-related proteins after TH5487 treatment revealed down regulation of Arg1, Mcp1 and Ccl11, and upregulation of the negative regulator of T_H_2, Bcl6. In addition, the OVA-induced airway hyperresponsiveness was significantly reduced by TH5487 treatment. Taken together, the data presented in this study suggest a clinically relevant utilization of TH5487 for the treatment of allergic inflammation.

**Graphical abstract:** 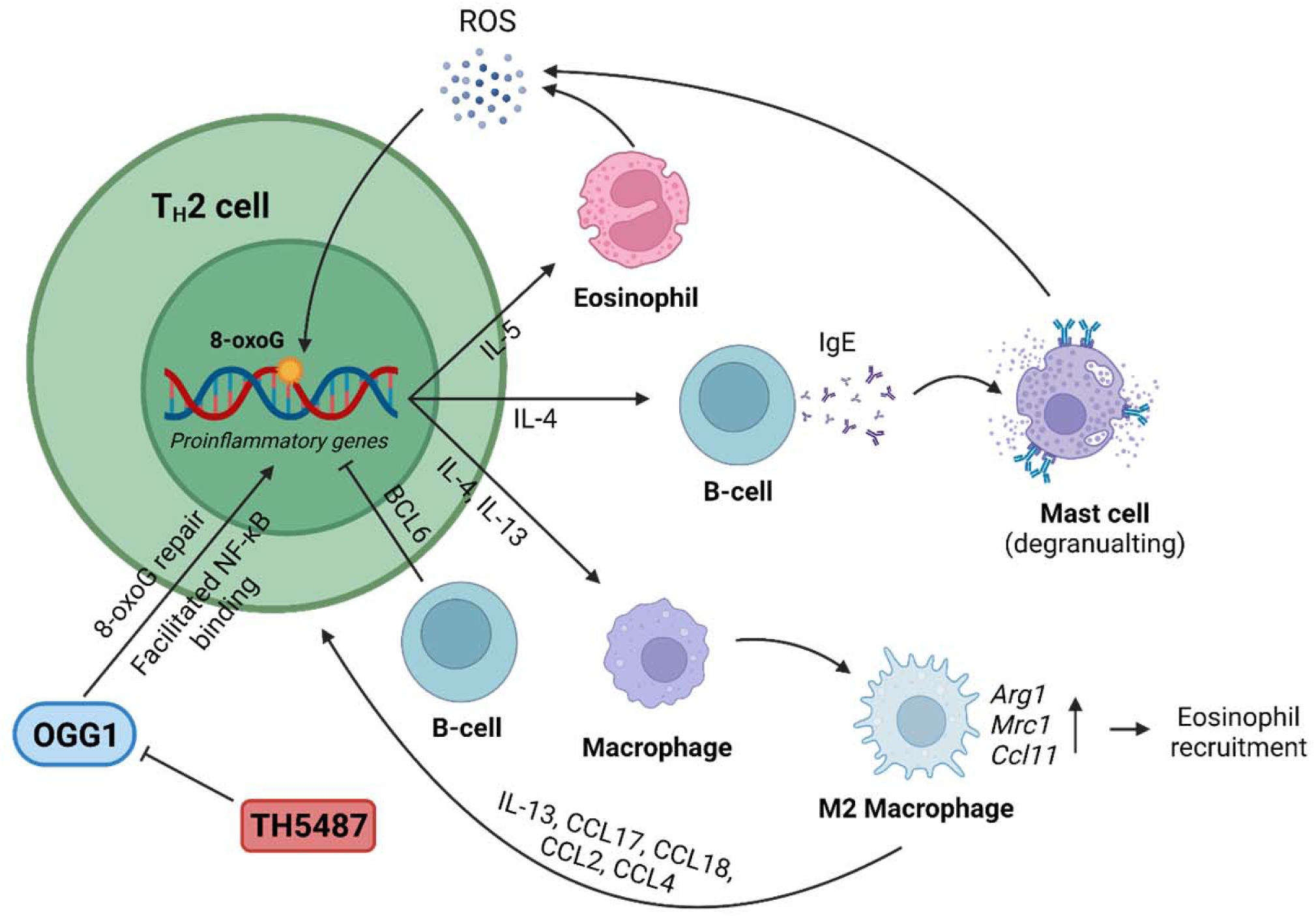

## Introduction

Asthma is a chronic inflammatory lung disease, affecting over 300 million people worldwide (The Global Asthma Report 2018, 2018). The disease is markedly heterogenous, complex, and partially heritable which is driven by environmental exposures (Hay, 2017). In sensitized individuals, disease is initiated by allergenic substances triggering expression of epithelial cytokines, the primary inducers of type 2 immune response to allergens (van Ree *et al*, 2014). Allergic asthma is an inflammatory disease characterized by increased mucus production, allergen specific IgE secretion, and accumulation of inflammatory cells, which leads to dyspnea, coughing, chest tightness, and airway remodeling (Rodrigo *et al*, 2004). Primary treatment options for asthma focus on reductions in lung inflammation using inhaled beta-agonists and corticosteroids (Reddel *et al*, 2021). However, a large proportion of asthma patients experience side effects, resulting in compromised treatment adherence (Cooper *et al*, 2015).

T helper 2 (T_H_2) lymphocyte driven airway inflammation is a key feature of allergic asthma, triggered by environmental antigens found in pollens, dust mites, fungi and pet dander (Permaul *et al*, 2012; Sporik *et al*, 1990; Suphioglu *et al*, 1992). Release of T_H_2 cytokines, including IL-4, IL-5 and IL-13, results in an infiltration of several different immune cells mainly comprising eosinophils, with the inclusion of mast cells, macrophages, monocytes, basophils, and neutrophils (Djukanovic *et al*, 1990; Holgate, 2012). IL-4 is key for the conversion of naïve helper T cells into the T_H_2 effector cells that promote IgE production, which binds to the surface of mast cells causing degranulation (Li-Weber & Krammer, 2003). IgE-mediated degranulation of mast cells leads to release of proinflammatory and cytotoxic mediators that increase lung inflammation (McBrien & Menzies-Gow, 2017; Bradding *et al*, 2006). IL-5 is also an important regulator of eosinophils, which play a vital role in airway remodeling during allergic asthma, controlling differentiation, activation, and delaying apoptosis of immune cells (Takatsu & Nakajima, 2008).

Oxidative stress in the airways occurs during asthma due to release of reactive oxygen species (ROS) from activated inflammatory cells, in particular eosinophils (Kirkham & Rahman, 2006). Increased ROS levels in the lungs of asthmatic patients are considered key factors in the pathogenesis of disease processes generated by the inhalation of environmental factors (Kirkham & Barnes, 2013). One of the most abundant DNA lesions is 8-hydroxy-2′-deoxyguanosine (8-oxoG), due to guanine possessing the lowest oxidation potential amongst DNA bases (Wu *et al*, 2004). Increased levels of oxidative damage to DNA in the form of 8-oxoG has been found in asthma patients (Proklou *et al*, 2013; Zeyrek *et al*, 2009). 8-oxoG lesions in chromatin of eukaryotic cells are predominantly repaired through base excision repair, which is initiated by 8-oxoG DNA glycosylase (OGG1) (David *et al*, 2007; Ba & Boldogh, 2018a). Recent studies have also documented 8-oxoG formed in promoter-enhancer regions, with OGG1 shown to be a modulator of gene expression via the facilitation of transcription factor DNA occupancy(Aguilera-Aguirre *et al*, 2017; Fleming & Burrows, 2017, 2020; Fleming *et al*, 2017; Li *et al*, 2012a; Tumurkhuu *et al*, 2020; Ba & Boldogh, 2018b). *Ogg1*^-/-^ murine studies (Klungland *et al*, 1999) highlighted resistance against LPS-induced inflammation including a reduced allergic inflammatory response after ovalbumin (OVA) challenge in OVA-sensitized mice (Mabley *et al*, 2005). To further investigate the role of OGG1 in allergic airway inflammation, OGG1 was depleted from the airway epithelium using siRNA in ragweed pollen grain extract-sensitized mice (Bacsi *et al*, 2013). Subsequent challenging with pollen extract intranasally resulted in a significantly lower allergic inflammatory response in the OGG1 siRNA treated mice compared to the mice receiving control siRNA. Moreover, *Ogg1^-/-^* mice exhibited metabolic disorders (Komakula *et al*, 2018; Sampath & Lloyd, 2019; Sampath *et al*, 2012a; Burchat *et al*, 2021), which has been shown to be an important comorbidity with asthma (Peters *et al*, 2018; Miethe *et al*, 2020; Mohan *et al*, 2019).

Recently, a small molecule inhibitor of OGG1, TH5487, was developed and shown to interfere with the binding of OGG1 to DNA in guanine-rich promotor regions, leading to reduced immune cell recruitment (Visnes *et al*, 2018). In this study, we assessed the potential therapeutic use of TH5487 in a mouse model of allergen-induced airway inflammation. TH5487 treatment resulted in reduced immune cell recruitment to the lungs, lower levels of plasma IgE and OVA-specific IgE, decreased NF-κB activation in the lungs, reduced small air way mucus accumulation and less M2 macrophages in BALF and lung tissue. Together, these results suggest a potential role for TH5487 as a treatment for allergic asthma.

## Methods

### Study design

Methods not described here can be found in the supplementary material. The aim of this study was to evaluate the treatment potential of TH5487 against allergic asthma using an OVA-induced allergic inflammation mouse model, which is a well-established model of allergic asthma. The mice were randomly divided into three equal groups with 5 animals in each group. When possible, downstream experiments were conducted with the investigator blinded to the sample groups. No animals were excluded as outliers and all experiments were performed in 2-4 technical replicates.

### Ethical approval

Animal experiments were approved by the Malmö-Lund Animal Care Ethics Committee, ethical permit no. M3802-19 and Stockholm Animal Care Ethics Committee, ethical permit no. 3649-2019.

### Animals

Female C57BL/6J mice (Janvier, Le Genest-Saint-Isle, France) and male BALB/c (Envigo, Horst, NL) 8-10-week-old mice were housed in plastic cages with absorbent bedding material and were maintained for at least 2 weeks prior to initiation of experiments. The mice were kept in a controlled environment (temperature, light/dark cycle, food and water *ad libitum*). Allergic airway inflammation was induced by sensitization with 20 µg OVA in alum (1:10) injected intraperitoneally at day 0 and 7 followed by challenges using intratracheal administration of 50 µg OVA at day 14, 16, 18 and 20 (Fig. 1A). An intraperitoneal injection of TH5487 (40 mg/kg) was performed prior to each challenge and mice were sacrificed at day 21. The mice were randomly allocated into three groups: OVA sensitized (vehicle), OVA sensitized + OVA challenged (OVA), and OVA sensitized + OVA challenged + TH5487 (OVA/TH5487). The lung function experiments were performed in BALB/C mice as this strain develops a stronger airway hyperresponsiveness (AHR) than C57BL/6 mice (Swedin *et al*, 2010).

**Figure 1:**
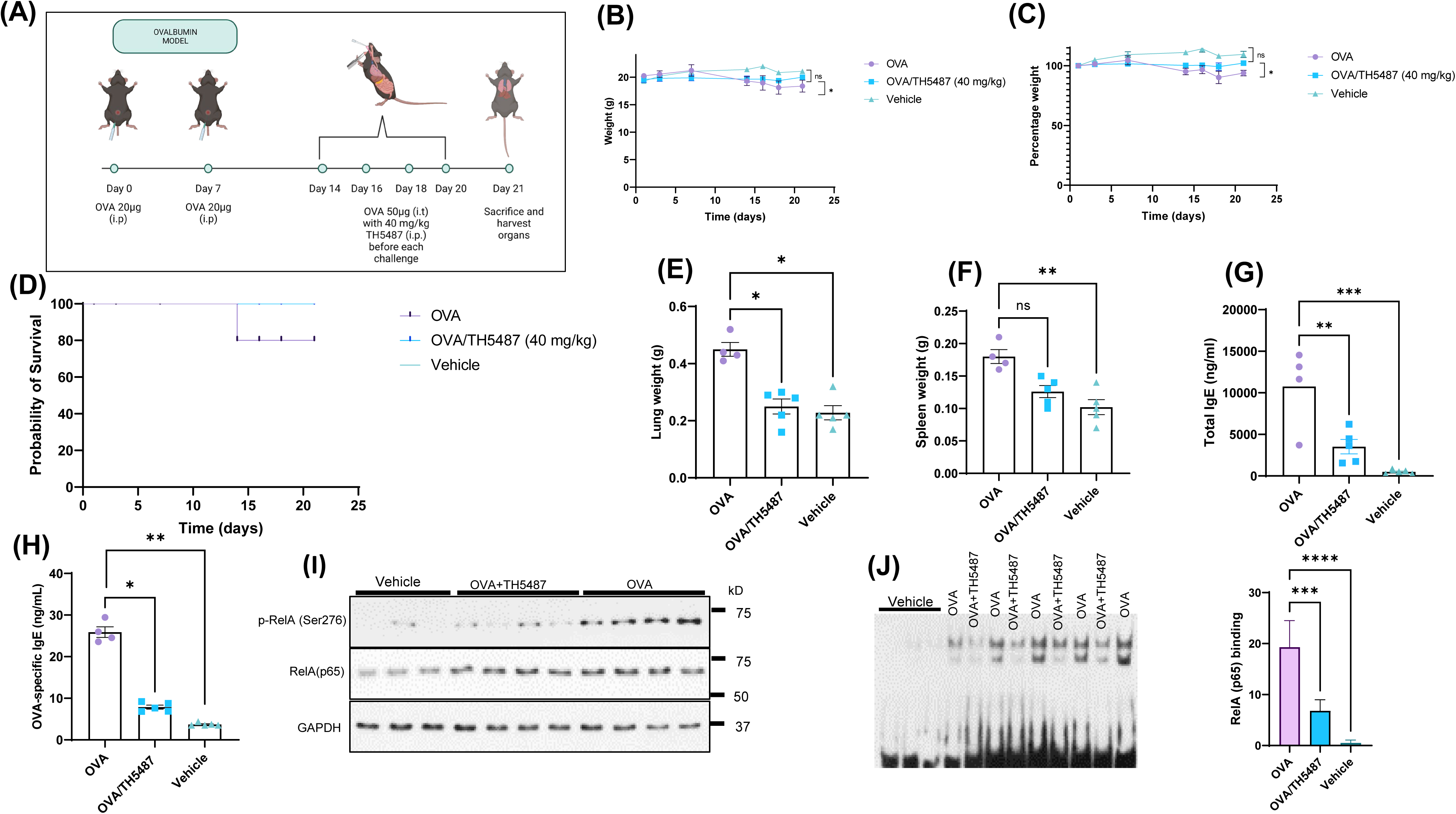
The effect of TH5487 treatment on survival, organ weights, NF-κB activation and IgE levels. (A) Sensitization and experimental schedule. All mice were sensitized with 20 µg ovalbumin (OVA) intraperitoneally at day 0 and 7. Mice in the OVA and OVA/TH5487 groups were challenged by intratracheal administration of 50 µg OVA at day 14, 16, 18 and 20. TH5487 treatment was performed by an intraperitoneal injection of 40 mg/kg before each challenge and all mice were sacrificed at day 21. (B and C) Total weights remained stable in all groups over the 21 days. (D) Probability of survival showing insignificant differences between the groups. (E) Lung weight comparison between the groups showing a significant increase in the OVA group compared to OVA/TH5487 and vehicle. (F) Spleen weights showing a significant increase in the OVA group compared to vehicle and a trend towards an increase compared to TH5487. (G) Total and OVA-specific (H) IgE levels in plasma, which was increased in the OVA group and significantly reduced after TH5487 treatment. (I) Western blot analysis of RelA and phosphorylated RelA(NF-κB) (p-p65/RelA) showing a significant decrease in OVA/TH5487 compared to OVA. (J) Binding of NF-κB homo (p50-p50) and heterodimer (p50-p65) to consensus DNA sequences (5’-GGGRNYYYCC-3’) in extracts derived from individual lungs of control (vehicle), OVA/TH5487 and OVA-challenged mice. Statistical comparison between the groups were performed using a one-way ANOVA followed by a Dunnett’s post-hoc test (\*\*\**P*<0.001, \*\**P*<0.01, \**P*<0.05).

### Blood collection

Collection of blood in tubes containing 0.5 M EDTA was performed by cardiac puncture. The tubes were centrifuged at 1,000 x *g* for 10 min and the supernatants were kept for later analysis.

### Lung tissue collection

The left lung from each mouse was collected in Histofix (Histolab, Göteborg, Sweden) and transferred to buffered paraformaldehyde solution (4%). The right lung was submerged in RNAlater solution (Thermo Fisher Scientific, Waltham, MA) and stored at -20 °C. After thawing, sample aliquots were homogenized in tissue protein extraction reagent (T-PER) solution (Thermo Fisher Scientific) with proteinase inhibitor (Pefabloc, SC; Sigma-Aldrich, Saint Louis, MI) at a concentration of 1 mM. Following homogenization, samples were centrifuged at 9,000 x *g* for 10 min at 4 °C and the supernatants were collected for later analysis. The remaining sample was used for RNA extraction (see below).

### Bronchoalveolar lavage fluid (BALF) collection

BAL was performed using a total volume of 1 ml PBS with 100 µM EDTA. The BALF was kept on ice and aliquoted for flow cytometry, cytospin differential counts and multiplex cytokine analysis. Cytospin samples were stained with modified Wright-Giemsa stain (Sigma-Aldrich) or used for immunofluorescence staining (see below).

### SDS-PAGE and western blotting

Total protein concentrations of lung homogenate lysates were analyzed with a Pierce BCA Protein Assay Kit (Thermo Fisher Scientific). SDS-PAGE was performed using Mini-PROTEAN^®^ Precast Mini PAGE Gels (Bio-Rad, Hercules, CA). Trans-Blot Turbo Mini 0.2 µM PVDF Transfer Packs (Bio-Rad) were used for transferring of proteins to the PVDF membranes. Membranes were blocked for 3h at RT and incubated with primary antibodies (rabbit anti-mouse NF-κB(RelA/p65/) (sc-8008; Santa Cruz Biotechnology, Santa Cruz, CA), rabbit anti-mouse phospho(p)-RelA/p65(NF-κB) (Ser276; A1953; Abcam), mouse anti-mouse arginase-1 (ab239731), rabbit anti-mouse CD206 (ab64693) and rabbit anti-mouse GAPDH (14C10; Cell Signaling; 1:500) in blocking buffer. After washing with PBS-Tween, membranes were incubated with secondary antibodies (Alexa Fluor 488-conjugated goat anti rabbit/mouse (Invitrogen, Carlsbad, CA) for 1h RT. Imaging of blots were preformed using a ChemiDoc system (Bio-Rad) followed by quantification with densitometry normalized to GAPDH.

### Total IgE ELISA

Total plasma IgE levels were determined using a IgE Mouse ELISA Kit (Invitrogen, Waltham, MA). Plasma samples diluted 1:500, and IgE standard (0.137-100 ng/ml) were pipetted into a 96-well plate and incubated for 2,5 h (RT) with gentle shaking. Wells were washed 4 times with wash buffer followed by addition of biotin conjugate to all wells. After 1 h incubation at RT with gentle shaking, the wells were washed 4 times and Streptavidin-HRP solution was added to all wells. The wash was repeated and TMB substrate was added following incubation for 30 min at RT with gentle shaking. The reaction was stopped by adding Stop Solution and absorbance was read at 450 nm using a VICTOR 1420 Multilabel plate reader (PerkinElmer, Waltham, MA). A standard curve was generated (Sigmoidal, 4PL) and used to calculate IgE levels in the samples.

### OVA-specific IgE ELISA

OVA-specific IgE levels were measured using a LEGEND MAX™ Mouse OVA Specific IgE ELISA Kit (BioLegend, San Diego, CA). The plate was washed 4 times with wash buffer followed by addition of Matrix A to standard wells and Assay Buffer to sample wells. Standard (20-0.313 ng/ml) and samples (diluted 1:2) were added to the plate and incubated for 2 h with shaking at 200 rpm. After washing the wells 4 times, Mouse Detection Antibody solution was added to each well followed by incubation for 1 h with shaking at 200 rpm. Wells were washed 4 times and Avidin-HRP D solution was added to each well and incubated for 30 min while shaking. The wells were washed 5 times, with soaking of the wells for 30 s to 1 min between each wash, before adding Substrate Solution F followed by incubation for 15 min in the dark. The reaction was stopped with Stop Solution and the absorbance was read at 450 nm using a VICTOR 1420 Multilabel plate reader (PerkinElmer). Using the absorbance of the standard, a linear standard curve was generated to calculate IgE levels in the samples.

### MUC5AC ELISA

The standards and BAL fluid samples were added to plates pre-coated with a MUC5AC antibody (MyBioSource Cat # MBS2507150). Thereafter biotinylated antibody specific for MUC5AC and then Avidin-Horseradish Peroxidase (HRP) conjugate was added to each well. Plates were washed and substrate added to each well. Reactions were terminated by the addition of a sulphuric acid solution. The optical density was measured spectrophotometrically at a wavelength of 450 nm using Synergy H1 BIO-TEK Instruments (Winooski, VT, USA).

### Immunostaining of lung sections

Fixated lung tissue sections underwent antigen retrieval (pH 9 buffer) using a Dako PT Link pre-treatment module (Agilent, Santa Clara, CA). Samples were washed with PBS and blocked with Dako protein block (Agilent) for 10 min, followed by incubation with primary antibodies overnight (mouse anti-mouse arginase-1 (ab239731), mouse anti-mouse MUC5ac (MA5-12178)). Samples were incubated with secondary antibodies, Alexa Fluor 594 goat anti mouse (Abcam, Cambridge, UK). Glass cover slips were mounted with DAPI-containing fluoroshield (Abcam). Images were visualized using a Nikon Confocal Microscope (Nikon, Tokyo, Japan) and fluorescence was quantified using ImageJ software (https://imagej.nih.gov).

### Immunostaining of BALF samples

Cytospin preparations of BALF cells were blocked with Dako protein block (Agilent) for 10 min, and incubated with primary antibodies (mouse anti-mouse arginase-1 (ab239731)) for 1 h RT. Secondary antibodies, Alexa Fluor 594 goat anti-mouse (Abcam), was added to the samples, followed by incubation for 30 min at RT. Slides were mounted with cover slips and DAPI-containing fluoroshield (Abcam). Images were visualized using a Nikon Confocal Microscope and fluorescence was quantified using ImageJ software.

### Real-time PCR Array

Total mRNA was extracted from lung tissue submerged in RNAlater using a RNeasy Mini Kit (Qiagen, Hilden, Germany) according to the protocol from the manufacturer. RNA concentrations were determined using a NanoDrop ND1000 (Saveen Werner, Limhamn, Malmö). Equal amounts of RNA were pooled from all animals in each group (Vehicle n=5, OVA/TH5487 n=5, OVA n=4). cDNA was synthesized with an iScript Advanced cDNA Synthesis Kit (Bio-Rad) and mixed with RT^2^ SYBR® Green ROX™ qPCR Mastermix. A volume of 25 µl of the reaction mixture was added to each well of a RT^2^ Profiler™ PCR Array Mouse Allergy & Asthma PAMM-067ZA plate. The RT-PCR reaction was performed using a QuantStudio™ 7 Flex system (Thermo Fisher Scientific) and data analysis was performed using the manufacturer’s web-based software (https://geneglobe.qiagen.com/analyze). Normalization of gene expression was performed using the following house-keeping genes: *B2m*, *Actb*, *Gusb*, *Gapdh* and *Hsp90ab1*.

### Electrophoretic mobility shift assay (EMSA)

Snap frozen lungs were homogenized and nuclear extracts were obtained using the CelLyticTM NuCLEARTM Extraction Kit (Millipore-Sigma). Protein concentrations were determined by Pierce BCA Protein Assay Kit (Thermo Fisher Scientific). EMSA assays were performed as described previously (Hao *et al*, 2018; Pan *et al*, 2016). Briefly, biotin-labeled probes (20 fmol; Sense: 5’-TTCCCTGGTCCCCGGGCTTTTCCAGACATCG-3′Anti-sense:5’- biotin CGATGTCTGGAA AAGCCCGGGGACCAGGGAA-3’) were mixed with 2 µg extract in binding buffer (10 mM Tris-Cl (pH 8.0), 10 mM NaCl, 1 mM DTT, 1 mM EDTA, 1 mg/ml BSA and 0.1 µg/µl Poly[d(I-C). Protein-DNA complexes were resolved on 6% nondenaturing polyacrylamide gels (Invitrogen) in 0.25 × TBE buffer (100V for 1.5h). Images were visualized using Amersham Imager 680 (Global Life Sci. Sol. Marlborough, MA). Band intensities were quantified using Image J v1.51 (U. S. NIH, Bethesda, Maryland, USA).

### Flow cytometry

A BD Accuri 6 (BD, Franklin Lakes, NJ) was used for the flow cytometry experiments. Cells were washed and stained with Fixable Viability Stain 510 (FVS510) (BD564406). The cells were washed in stain buffer 1x (BD554656) followed by incubation with Lyse Fix 1x (BD558049 (5x)). After fixing the samples were washed with stain buffer and separated into two equal samples. The first sample was incubated with either anti-CD11b (BD553312), anti-CD11c (BD558079) and anti-Ly6G (BD551461), with the second sample incubated with anti-CD11b (BD553312), anti-CD11c (BD558079) and anti-SiglecF (BD562680).

### Bioplex cytokine analysis

A Bio-Plex Pro mouse cytokine assay (23-Plex Group I; BioRad) using a Luminex-xMAP/Bio-Plex 200 System was used to quantify multiple cytokines in BALF, plasma and lung homogenate. Analysis was performed using Bio-Plex Manager 6.2 software (Bio-Rad). The detection limits were as follows: Eotaxin (21372.02-1.15 pg/mL), GCSF (124018.4-6.97 pg/mL), GMCSF (1161.99-3.73), IFN-γ (14994.64-0.72 pg/mL), IL-1α (10337.5-0.63 pg/mL), IL-1β (28913.54-1.57 pg/mL), IL-2 (22304.34-1.21 pg/mL), IL-3 (7639.21-0.47 pg/mL), IL-4 (6334.86-0.36 pg/mL), IL-5 (12950.39-0.76 pg/mL), IL-6 (11370.16-0.66 pg/mL), IL-9 (2580.93-2.46 pg/mL), IL-10 (76949.87-4.09 pg/mL), IL-12p40 (323094.58-17.38 pg/mL), IL-12p70 (79308.46-19.51 pg/mL), IL-13 (257172.3-53.85 pg/mL), IL-17 (8355.61-0.5 pg/mL), KC (23377.88-1.3 pg/mL), MCP-1 (223776.6-45.04 pg/mL), MIP-1α (14038.07-0.58 pg/mL), MIP-1β (928.18-2.39 pg/mL), RANTES (4721.74-4.42 pg/mL), and TNF-α (73020.1-4.61 pg/mL). Correction for protein concentration was done using a Pierce ™ BCA Protein Assay Kit (Thermo Fischer Scientific).

### H&E and PAS staining of lung sections

Right lungs were fixed in Histofix (Histolab) and paraffin embedded. Sections (3 µm) were cut with a microtome and placed on glass slides (Superfrost Plus; Thermo Fisher Scientific). Deparaffinization was performed using serial baths of xylene and ethanol. Staining was completed using Mayer hematoxylin and 0.2 % eosin (Histolab) or Periodic Acid Schiff (PAS) Stain Kit (Mucin Stain) (Abcam). Imaging of the slides was performed using an Olympus BX60F microscope with an SC50 camera (Olympus, Tokyo, Japan).

### Real-time PCR

Total mRNA was extracted from lung tissue kept in RNAlater stored at -20 °C using a RNeasy Mini Kit (Qiagen, Valencia, CA) according to the manufacturer’s protocol. RNA concentrations were determined with a NanoDrop ND1000 (Saveen Werner, Malmö, Sweden). RNA was converted into cDNA (1 µg) using an iScript Advanced cDNA Synthesis Kit (Bio-Rad). Expression of target genes was measured using TaqMan™ Fast Advanced Master Mix with TaqMan™ probes listed in **Supp. Table 1**, and the reactions were run on a CFX Connect Real-Time System in 96-well plates. Water samples were included to confirm non-specific PCR reactions. ΔCt values were calculated by normalization to house-keeping gene succinate dehydrogenase complex, subunit A (*Sdha*). To obtain ΔΔCt values, ΔCt values from the treatment groups were divided by ΔCt values from the control group. Fold change was calculated by 2^-ΔΔCt^.

### Measurement of Airway hyperresponsiveness (AHR)

On day 21 of the OVA-challenge protocol, AHR was induced by administration of methacholine (MCh; Sigma-Aldrich) in mice anaesthetized with ketamine hydrochloride (75mg/kg, Ketaminol® Vet., Intervet, Stockholm, Sweden) and medetomidine hydrochloride (1mg/kg, Cepetor®Vet., VETMEDIC, Stockholm, Sweden). Methacholine was delivered by aerosol administration via a nebuliser (Scireq; Montreal, Que., Canada), at doses ranging from 0 to 12.5 mg/ml after an initial dose of saline alone. The AHR was measured with a small animal ventilator (FlexiVent; Scireq), as previously described. Dynamic and central resistance (R and Rn, respectively), central airway compliance (C), peripheral tissue damping (G) and tissue elastance (H) were recorded. Newtonian resistance, Rn, is a predictive measure of resistance in the central airways, with tissue damping reflecting energy dissipation in the lung tissue, and tissue elastance indicating tissue stiffness.

### Statistical analysis

Analysis of difference between three or more groups was calculated using one-way ANOVA with Dunnett’s post hoc test. Statistical testing was performed using GraphPad Prism 9.3.1 (350) (GraphPad Software, San Diego, CA) and the statistical significance was defined as *P* < 0.05. The results are displayed as mean ± SEM.

## Results

### TH5487 treatment decreases levels of activated NF-κB and plasma IgE

Allergic airway inflammation was induced by ovalbumin (OVA)-sensitization and challenge (**Fig. 1A**). An intraperitoneal injection of TH5487 (40 mg/kg) was performed prior to each challenge. No significant differences in weight loss or probability of survival were seen between the groups (**Fig. 1B-D**). Lung weights were significantly increased in the OVA group compared to the TH5487 treated mice (**Fig. 1E**) and a similar trend was seen for the spleen weight, without reaching statistical significance (**Fig. 1F**). Total IgE and OVA-specific IgE levels in plasma were both significantly decreased in the OVA/TH5487 group compared to the OVA group (**Fig. 1G** **and** **H**). Activation of the NF-κB signaling pathway plays a central role in allergic inflammation of both patient-derived samples and in OVA-sensitized mice (Gagliardo *et al*, 2003; Poynter *et al*, 2002). Western blot analysis of NF-κB displayed increases in the phosphorylated catalytic subunit (p-RelA (p65; the mammalian homolog of the V-Rel avian reticuloendotheliosis viral oncogene A) in the OVA-challenged group, whereas treatment with TH5487 resulted in no significant changes in levels of p-RelA/p65(NF-κB) (**Fig. 1I**). To support these observations, EMSAs were performed, showing increased binding to probes of homo (p50-p50) and heterodimeric (pRelA-p50) complexes of NF-κB in lung extracts from OVA-challenged mice. In lung extracts of OVA challenged TH5487-treated mice there were decreased levels of both p50-p50 and p50-p65 bound to probe (**Fig. 1J**).

### Goblet cell hyperplasia and airway mucin production are decreased following TH5487 treatment

Lungs of sensitized animals challenged with OVA, OVA and TH5487-or vehicle treatment were sectioned and stained with H&E or PAS stains. Histological analyses of H&E-stained sections showed increased cellularity, especially around primary and secondary bronchi and bronchioles in OVA-challenged groups. Significantly decreased levels of accumulated inflammatory cells were seen in lungs of TH5487 treated animals (**Fig. 2A** **and Supp. Fig. 1-3**). After PAS staining, bronchiolar mucosal epithelium in lungs of OVA-challenged animals exhibited increased epithelial cell hyperplasia (**Fig. 2B**). TH5487-treatment of OVA-challenged mice significantly (\**P* < 0.001) decreased PAS positive cells compared with OVA alone (**Fig. 2B**), while vehicle alone showed no PAS positive cells (**Fig. 2B**). **Fig. 2C** shows immunochemical staining of MUC5AC (upper panel) in OVA, which was decreased by TH5487 treatment of OVA sensitized/challenged animals. The histological observations are supported by ELISA of BALF, showing significantly decreased MUC5AC levels in the OVA/TH5487, compared to that of OVA alone group (**Fig. 2D**).

**Figure 2:**
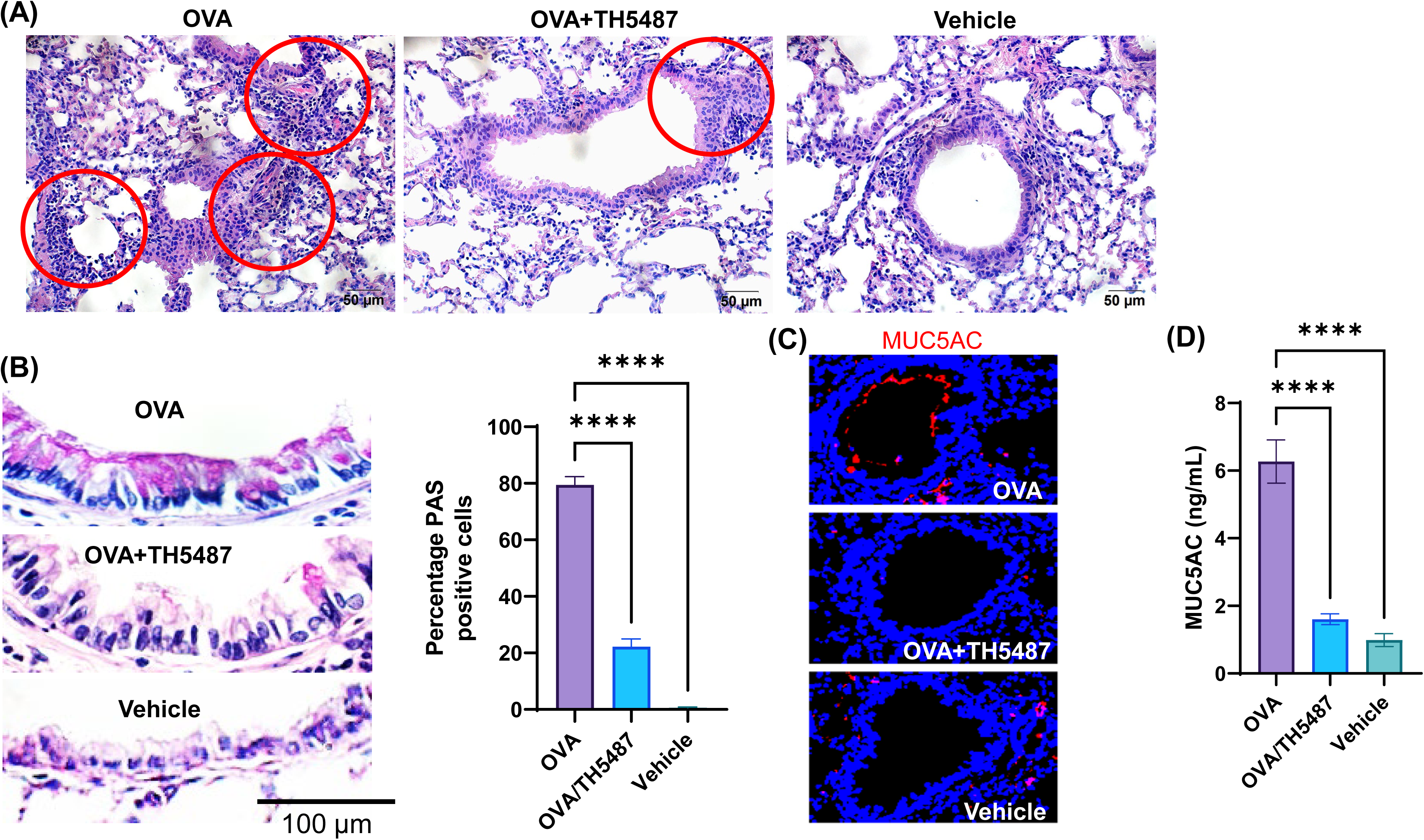
H&E, PAS and immunofluorescence staining of murine lung sections. Mouse lungs were harvested, formalin fixed, sectioned, and stained with haemotoxylin and eosin (H&E), Periodic acid-Schiff (PAS) or fluorescent antibodies. (**A**) Representative images of H&E-stained lung sections from OVA-challenged mice with and without TH5487 treatment. Inflammatory infiltrates surrounding bronchi/bronchioles (red circles); OVA: lung section of ovalbumin-challenged mouse; OVA + TH5487: lung section of TH5487-treated OVA challenged mouse; right panel, control vehicle alone. Scale bar: 50 µm. (**B**) TH5487 decreases mucin-containing cells (Goblet cells) as shown by PAS staining (magenta). Upper panel, a representative image from unchallenged (vehicle), OVA (middle) and OVA/TH5487 (lower panel) challenged/treated lungs. Magenta-colored epithelial cells are positive for mucin. Right panel, graphical depiction of PAS positive cells in airway epithelium. Percentage of mucin producing cells were enumerated in epithelium of bronchioles from 3 sections of each lung by two independent investigators. Percentage of mucin positive cells were calculated and graphically depicted. Scale bar: 100 µm. (C) Murine lung sections were stained with MUC5AC antibody (red) to determine mucin production, with (D) MUC5AC ELISA conducted on homogenized murine lung tissues. Statistical comparison between the groups were performed using a one-way ANOVA followed by a Dunnett’s post-hoc test (\*\*\**P*<0.001, \*\**P*<0.01, \**P*<0.05).

### Immune cell recruitment to the lung is mitigated by treatment with TH5487

In allergic asthma, recruitment of immune cells to the airways results from an increased release of cytokines/chemokines from the airway epithelium and from resident immune cells (Velazquez & Teran, 2011). The effect of TH5487 on immune cell recruitment to the lungs was investigated by performing flow cytometry on BALF from OVA-challenged mice, with the gating strategy shown in **Fig. 3A**. Large increases in the number of eosinophils, inflammatory macrophages, alveolar macrophages and neutrophils were seen in the OVA challenged mice compared to the control mice (**Fig. 3B** **and** **C**). Administration of TH5487 resulted in significant reductions of all immune cells, with almost no detection of eosinophils and macrophages. BALF samples stained with Giemsa-Wright revealed an increased number of immune cells in the OVA challenged mice compared to the TH5487 and vehicle treated mice (**Fig. 3D**).

**Figure 3:**
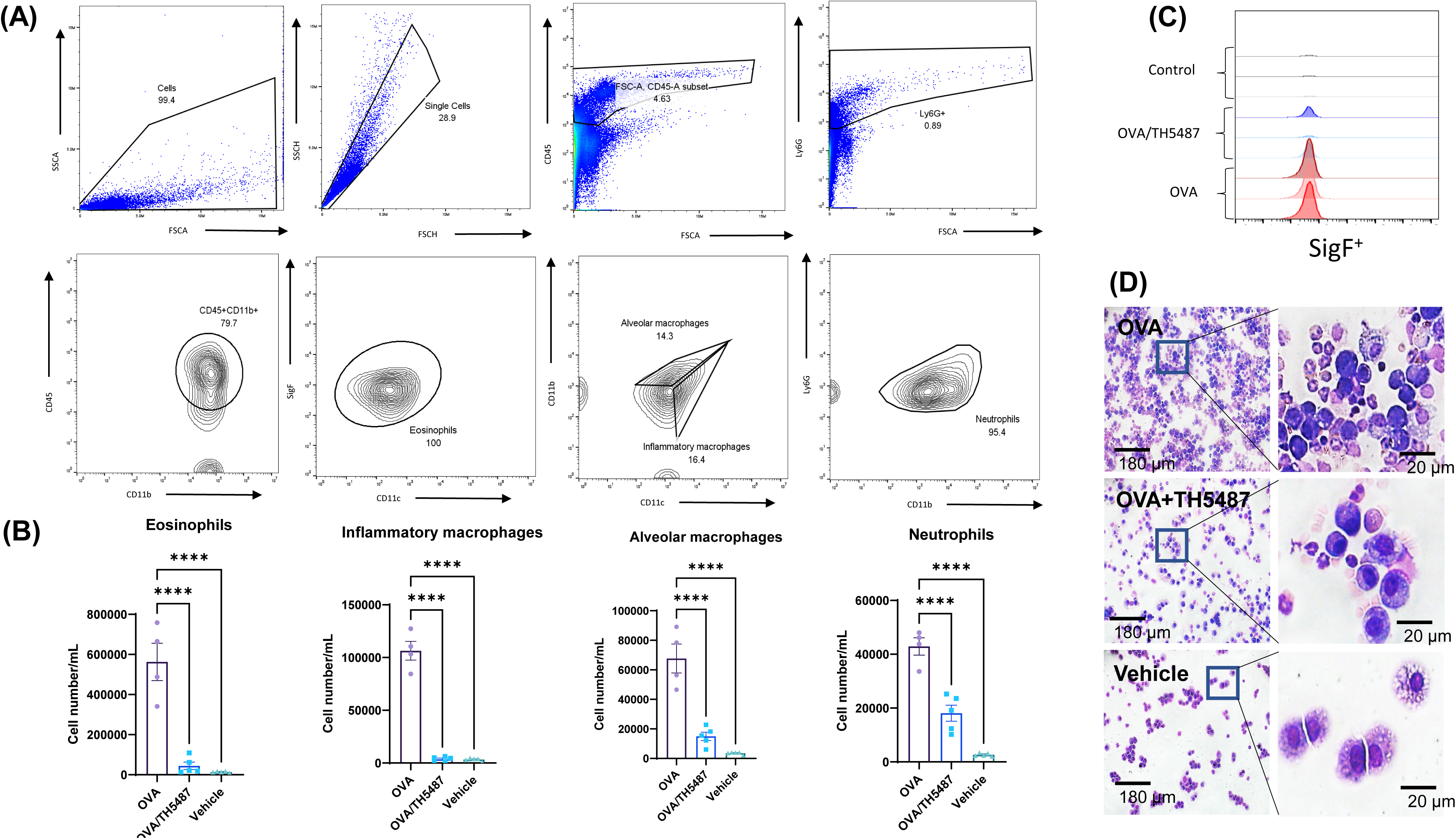
Immune cell recruitment measured in murine BALF. (A) Gating strategy used to determine eosinophils, inflammatory macrophages, alveolar macrophages and neutrophils in BALF samples. (B) A significant increase of eosinophils, inflammatory macrophages, alveolar macrophages and neutrophils was seen in the OVA-challenged group. TH5487 treatment significantly reduced recruitment of all cell types to the lung, with overlay histogram showing Siglec-F+/CD11b+ eosinophils. (D) Giemsa-Wright stained cytospins of BALF samples showing an increase in the number of immune cells in the OVA group compared to OVA/TH5487 and vehicle. Statistical comparison between the groups were performed using a one-way ANOVA followed by a Dunnett’s post-hoc test (\*\*\*\**P*<0.0001).

### Administration of TH5487 reduces murine cytokine levels in BALF, lung and plasma

Inflammatory cytokines were measured in lung homogenate, BALF and plasma using a 23-cytokine multiplex assay. Noticeable increases in most cytokines were seen in the OVA group, indicating increased allergic inflammation in the lungs of these mice (**Fig. 4 A-C, left and right panels and Supp. Fig. 4-6**). Treatment with TH5487 resulted in a significant decrease in a large majority of cytokines in BALF and lung homogenate. Moreover, a considerable decrease of the T_H_2 cytokines IL-4, IL-5 and IL-13 was shown in the TH5487 treated mice, suggesting a reduced T_H_2 response. Eotaxin (eosinophil chemotactic protein or C-C Motif Chemokine Ligand 11) levels in the BALF were significantly decreased after TH5487 treatment (*P*=0.0136), which partly explains the decreased recruitment of eosinophils to the lungs. A significant increase was also seen in monocyte chemotactic and activating factor (MCP-1 or CCL2) in the BALF (*P*=0.0174), indicating an increased recruitment of monocytes and macrophages to the lung. However, treatment with TH5487 resulted in a significant reduction of MCP-1 (*P*=0.0216) to levels nearly equal to the vehicle group, further supporting decreased immune cell recruitment to the lung.

**Figure 4:**
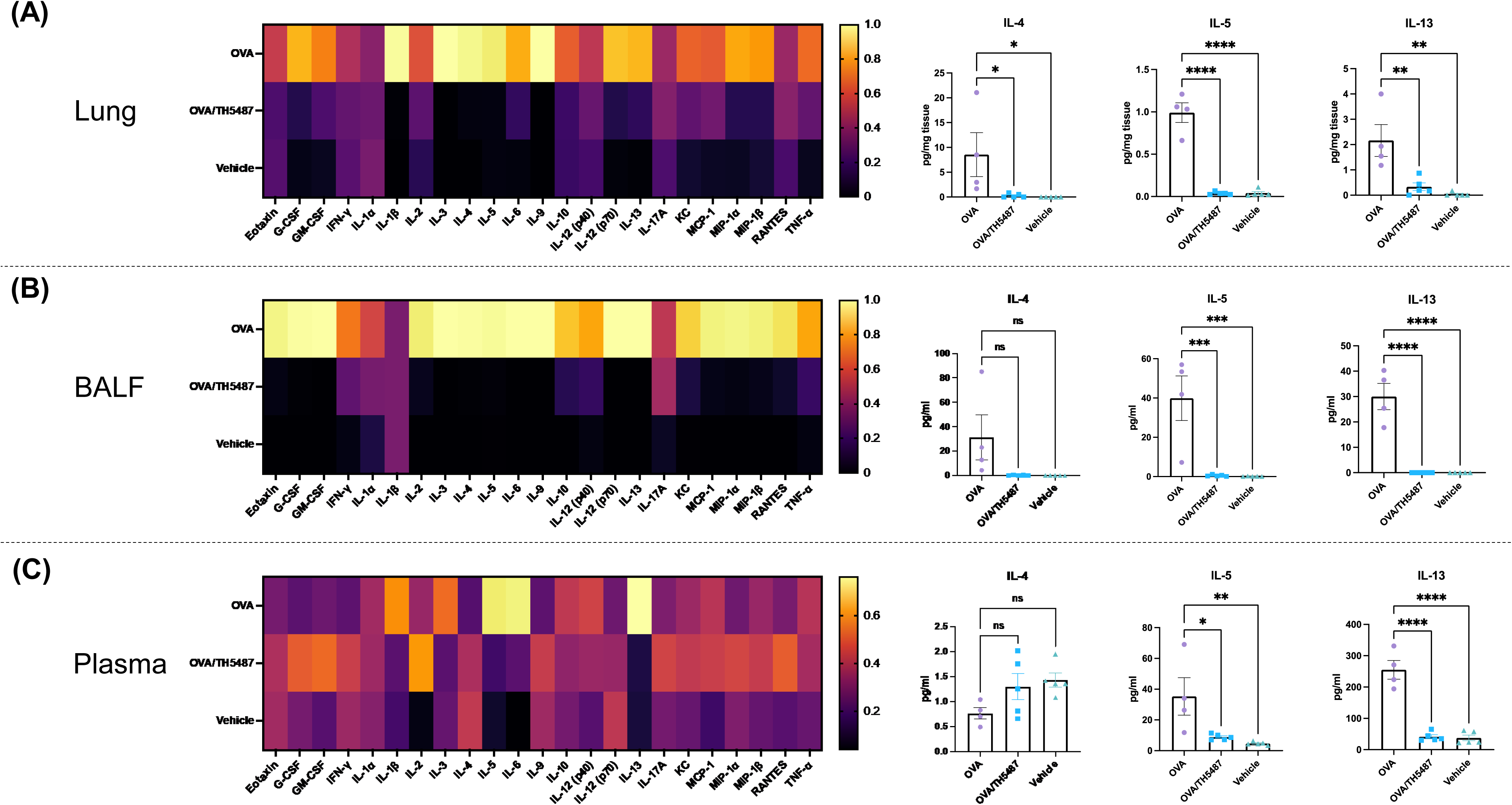
Murine cytokine levels in lung, BALF and plasma. (A-C) Heatmaps displaying differences in cytokine levels in lung homogenate, BALF and plasma (yellow indicates high value; black indicates low value). The key T_H_2 cytokines IL-4, IL-5 and IL-13 are shown as individual graphs including statistical comparisons. Differences in cytokine levels were compared using a one-way ANOVA followed by a Dunnett’s post-hoc test (\*\*\*\**P*<0.0001, \*\*\**P*<0.001, \*\**P*<0.01, \**P*<0.05).

### Gene expression profiling reveals decreases in expression of key T_H_2 response genes by TH5487 administration

Using a RT^2^ Profiler™ PCR Array Mouse Allergy & Asthma panel, the expression of 84 allergy and asthma related genes in the lung was quantified (**Fig 5A**). Comparison between the OVA and OVA/TH5487 groups revealed a difference in mRNA fold regulation of several genes (**Fig. 5B**). The largest decrease in expression (−10.99 fold) was seen in the macrophage M2-marker *Arg1*, indicating a decreased number of M2 macrophages in the lung. Furthermore, the expression of *Tnfrsf4*, which is involved in activation of T-cells, was decreased after TH5487 treatment (−4.76 fold). A decrease was also seen in CCL11 (−3.88 fold), which encodes the eosinophil specific chemotactic factor eotaxin-1. Two genes were shown to be upregulated, i.e. Bcl6 (2.55 fold) and Clca1 (3.42). Interestingly, Bcl6 reduces T_H_2 responses through transcriptional repression of several key cytokines and chemokines (Arima *et al*, 2008). Clca1 is a regulator of mucus production in goblet cells, but studies describing its role in allergic asthma have been contradictory (Robichaud *et al*, 2005; Nakanishi *et al*, 2001). Using Metascape, a resource for analysis of system-level datasets (Zhou *et al*, 2019), gene ontology (GO) terms associated with the results from the gene expression profiling were generated (**Fig. 5C** **and** **D**). The most significant GO terms were cytokine-cytokine receptor interaction (Log10P (−14.8)), eosinophil chemotaxis (Log10P (−13.9)) and eosinophil migration (Log10P (−13.8)).

**Figure 5:**
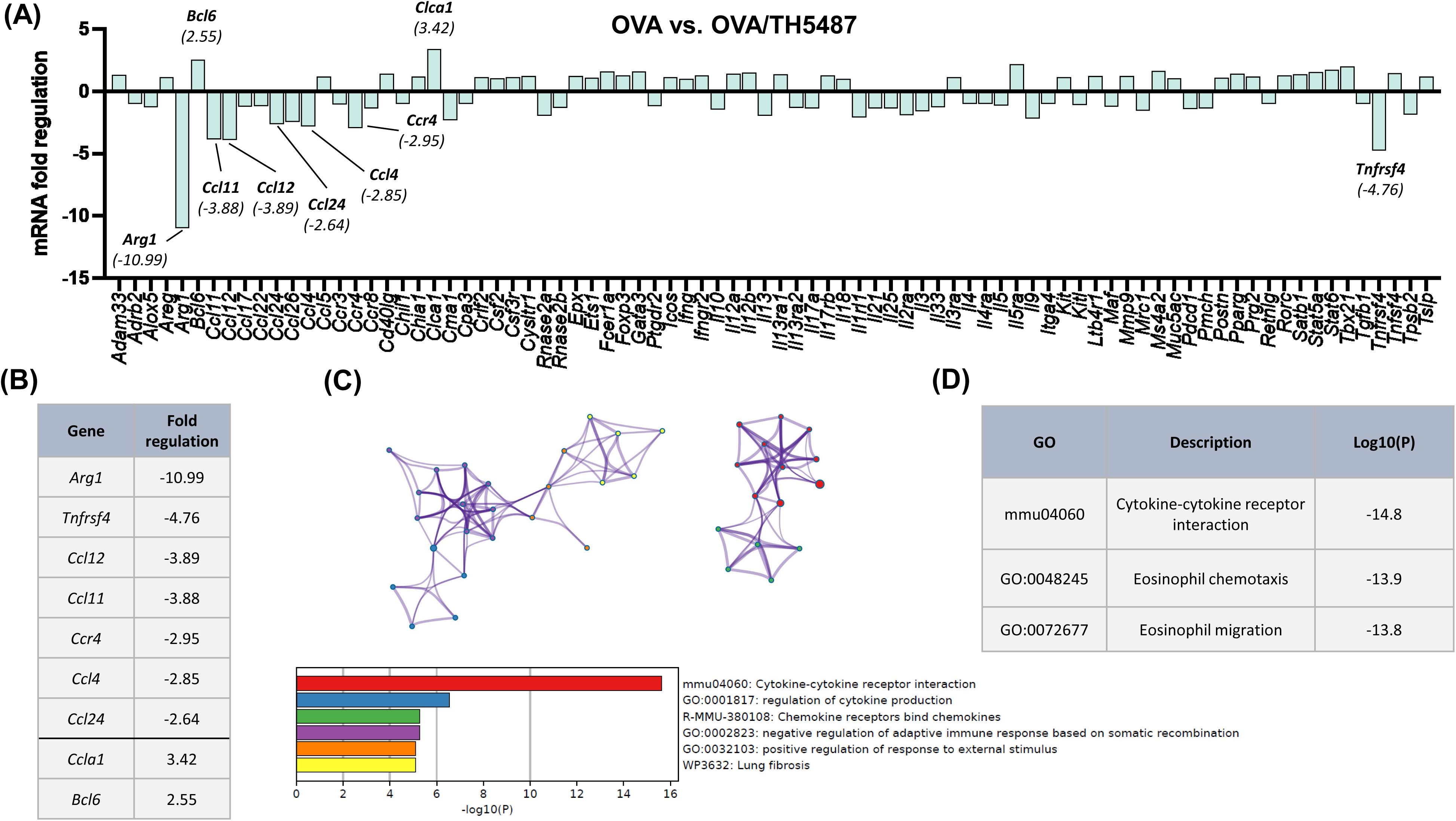
Asthma and allergy related gene expression profiling of lung tissue. (A) Asthma and allergy gene array comparison between OVA and OVA/TH5487. Equal amounts of mRNA from each mouse were pooled (OVA n=4, OVA/TH5487 n=5). Individual genes with a difference in fold regulation of >2.5 are highlighted in the graph. (B) List of genes with a fold regulation of >2.5. (C) Metascape protein network analysis displaying key proteins (>Log2.5) and the associated gene-ontology terms (color-coded). (D) Key gene-ontology terms displaying significantly downregulated Log10P values following treatment with TH5487.

### Macrophage M2 polarization is attenuated by TH5487 treatment

T_H_2 immune responses, which is a key feature of allergic asthma, has been shown to induce polarization of macrophages towards an M2 phenotype (Girodet *et al*, 2016; Nie *et al*, 2017). Gene expression of the M2 markers *Mrc1*, which encodes the surface receptor CD206, and *Arg1* in the lungs of the mice was shown to be significantly reduced in the OVA/TH5487 mice compared to the OVA/vehicle mice (**Fig. 6A**). Western blot analysis using lung homogenates revealed a significant increase in Arg1 in the OVA challenged mice compared to the TH5487 and vehicle treated mice (**Supp. Fig. 7A and 8**). A trend towards a similar difference was seen for CD206. Immunofluorescence staining of BALF samples with Arg1 antibodies showed a significant increase in Arg1 positive cells in the OVA group compared to OVA/TH5487 (**Supp. Fig. 7B**). Furthermore, Arg1 immunofluorescence staining of lung sections further confirmed a significant increase of Arg1 positive cells in the OVA group compared to the TH5487 treated group (**Fig. 6B**).

**Figure 6:**
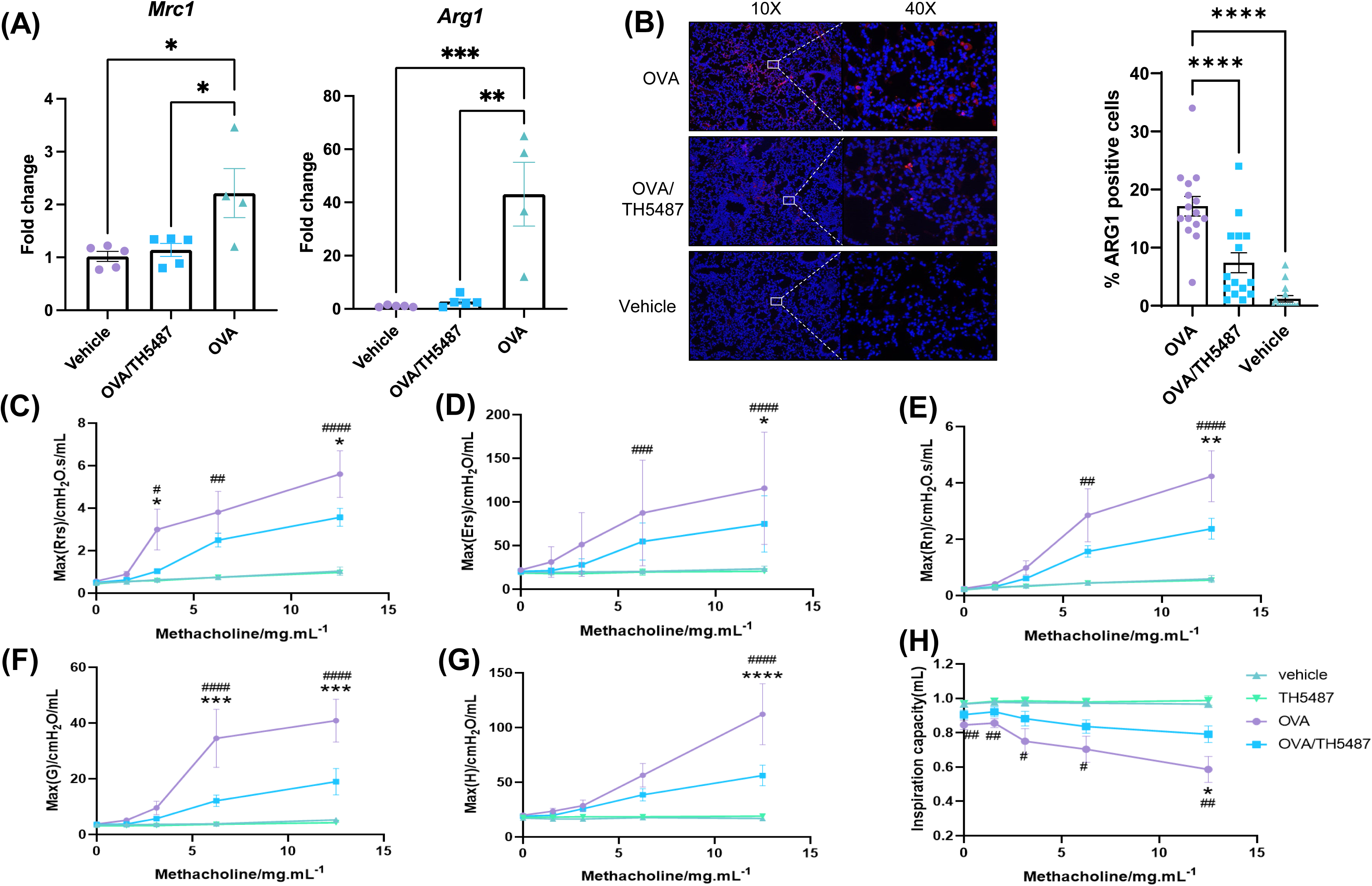
TH5487-mediated differences in macrophage M2 polarization and improved airway function in OVA-challenged mice. (A) RT-PCR measured expression of *Mrc1*, which encodes CD206, and *Arg1* is reduced in the OVA/TH5487 group compared to the OVA group. (B) Murine lung sections stained with Arg1 antibodies revealing decreases in Arg1 positive cells in the OVA/TH5487 group compared to OVA. Statistical comparisons were performed using a one-way ANOVA followed by a Dunnett’s post-hoc test (\*\*\*\**P*<0.0001, \*\*\**P*<0.001, \*\**P*<0.01, \**P*<0.05). The airway hyperresponsiveness (AHR) was measured with a small animal ventilator (FlexiVent; Scireq), as described in Materials and Methods. (C-E) Dynamic, elastic, and central resistance (Rrs, Ers, and Rn, respectively), peripheral tissue damping (F), tissue elastance (G), and central airway compliance (H) were recorded. The results are expressed as mean ± SEM (n = 4–9). Vehicle only (vehicle), TH5487 only (TH5487), ovalbumin-challenged (OVA), and ovalbumin-challenged/ TH5487-treated (OVA/TH5487). \**P*<0.05, \*\**P* <0.01, \*\*\**P*<0.001, \*\*\*\**P*< 0.0001 vs. respective OVA/TH5487, #*P*<0.05, ##*P* <0.01, ###*P*<0.001, ####*P*<0.0001 vs. respective vehicle.

### Improved lung function in OVA-challenged TH5487-treated animals

To assess AHR, OVA-challenged mice were administered methacholine (MCh), a substance known to induce smooth muscle constriction^52^ similar to the allergic asthma response seen clinically. Our results showed significantly elevated levels in OVA-challenged mice (*P*<0.05) of dynamic (Rrs) and central resistance (Rn) at 12.5 mg/mL MCh compared to the vehicle and TH5487-treated animals (**Fig. C-E**). TH5487/OVA mice displayed decreased tissue dampening and decreased tissue elastance compared to OVA-challenged mice (*P*<0.001 and *P*<0.0001, respectively; **Fig. 6F** **and** **G**). Subsequently, inspiration capacity was significantly decreased in OVA-challenged mice compared to TH5487-treated animals (*P*<0.05; **Fig. 6H**).

## Discussion

Allergic asthma is a highly complex and heterogeneous inflammatory disease involving multiple cell types and tissues (Holgate *et al*, 2015). Molecular mechanisms underlying its pathogenesis are not well understood, however, it is clear that ROS are key to allergic inflammation and oxidatively modified molecules including intrahelical 8-oxoG (Wu *et al*, 2000a; MacPherson *et al*, 2001a; Nadeem *et al*, 2003; Sahiner *et al*, 2011). Recent studies have identified 8-oxoG in promoter regions appearing as epitranscriptomic-like marks, with OGG1 DNA repair coupled to transcriptional regulation of chemokines and cytokines (Hao *et al*, 2020, 2018; Ba & Boldogh, 2018b; Ba *et al*, 2014). Therefore, we tested whether pharmacological targeting of OGG1 decreased allergic immune responses in a mouse model. We showed that the small molecule TH5487, which selectively inhibits the binding of OGG1 to oxidatively modified guanines, significantly decreased transcription from pro-inflammatory genes. Additionally, TH5487 treatment reduced the recruitment of eosinophils and other inflammatory cells and decreased mucin production, displaying promising results as potential treatment against allergic inflammation.

Allergic asthma is a disease characterized by dysregulated expression of T_H_2 cytokines in the airways, resulting in increased recruitment of inflammatory cells such as eosinophils and macrophages. Eosinophils are significant contributors to oxidative stress through the release of reactive oxygen species (ROS), including nitrogen oxide (NO)-derived oxidants that cause damage to tyrosine residues (MacPherson *et al*, 2001b; Wu *et al*, 2000b). CCL11 (eotaxin-1) plays a central role in the recruitment of eosinophils and is primarily produced by the epithelium in response to allergenic exposure within the airways. However, during an allergen challenge the primary source of CCL11 is switched to macrophages (Rothenberg, 1999). In asthmatic patients and OVA-sensitized mice, macrophages are polarized towards an M2 phentoype (Girodet *et al*, 2016; Nie *et al*, 2017), suggesting that M2 macrophages are an important source of CCL11 during allergic asthma. In this study, we found a reduced expression of CCL11 in lung tissue as well as a significant reduction of the levels in the BALF after treatment with TH5487, which could be explained by a reduction in M2 macrophages. Indeed, reductions in the expression of the two M2 markers *Arg1* and *Mrc1* were seen in lung tissue of mice treated with TH5487. This was further established by immunofluorescent staining of Arg1 in cytospins from BALF samples and lung sections, where TH5487 administration resulted in less immunoreactivity. M2 macrophages produce several different factors, including the anti-inflammatory cytokine IL-10. Studies in mice lacking expression of IL-10 have shown an increased survival of T_H_2 cells, leading to an increased expression of IL-5 which exacerbates pulmonary inflammation characterized by an influx of eosinophils (Yang *et al*, 2000; Coomes *et al*, 2017). Moreover, M2 macrophages further exacerbates the T_H_2 response by releasing IL-13, CCL17, CCL18, CCL22, CCL24 (eotaxin-2), and CCL26 (eotaxin-3), thereby increasing eosinophil chemotaxis (Siddiqui *et al*, 2013). Consequently, a reduction of M2 macrophages in allergic asthma would decrease the production of T_H_2 cytokines and reduce eosinophil recruitment to the lungs.

Gene expression profiling of lung tissue from OVA and OVA/TH5487 mice respectively, revealed differences in mRNA fold changes of several genes. Two genes were upregulated more than 2.5-fold, including Bcl6, which has been shown to negatively regulate T_H_2 responses and macrophage related chemokines (Arima *et al*, 2008). Knockout of Bcl6 in mice results in eosinophilic inflammation due to an overproduction of T_H_2 cytokines (Dent *et al*, 1997). The ability of Bcl6 to reduce the T_H_2 response is due to its ability to regulate the expression of several genes. Bcl6 decreases the expression of IL-5 in T-cells by binding to a silencing element of the IL-5 gene (Arima *et al*, 2002). Bcl6 also suppresses the expression of IL-6 and MCP-1 in macrophages (Yu *et al*, 2005; Toney *et al*, 2000) as well as MIP-1α and NF-κB in lymphocytes (Shaffer *et al*, 2000; Li *et al*, 2005). Finally, Bcl6^-/-^ mice have increased amounts of IgE due to immunoglobulin class switching in B-cells (Harris *et al*, 1999). In our study, administration of TH5487 resulted in significantly lower concentrations of IL-5, IL-6, MCP-1 and MIP-1α in BALF, decreased phosphorylation of NF-κB in the lungs and a reduction in total IgE and OVA-specific IgE, which is consistent with the increased expression of *Bcl6* after TH5487 treatment.

The effectiveness of TH5487 stems from the inhibition of OGG1, which has pleiotropic roles in gene expression by binding to intrahelical 8-oxoG located in regulatory regions and facilitates transcription factor’s DNA occupancy including NF-κB (Ghosh & Mitchell, 1999; Pan *et al*, 2017; Schreck *et al*, 1990; Hao *et al*, 2018). OGG1 creates specific DNA structural changes that facilitate NF-κB (and other transcription factors) recognition and binding to its consensus motif (Bruner *et al*, 2000; Ghosh & Mitchell, 1999; Pan *et al*, 2017; Schreck *et al*, 1990). In addition, OGG1-mediated incision of DNA as part of BER is also linked to modulation of gene expression (Pastukh *et al*, 2015; An *et al*, 2016; Zhu *et al*, 2018; Fleming *et al*, 2019). Excision of 8-oxoG by OGG1 generates apurinic sites in gene regulatory regions, particularly in promoters with potential G-quadruplex, Z DNA or i-motif-forming sequences often present in promoters of inflammatory genes (An *et al*, 2016; Zhu *et al*, 2018; Fleming *et al*, 2019). Studies using *Ogg1* knockout animals and biochemical approaches have shown distinct roles for OGG1 in gene expression in a stimuli and context-dependent manner (Hao *et al*, 2018; Perillo *et al*, 2008; Sampath & Lloyd, 2019; Sampath *et al*, 2012b; Simon *et al*, 2020). This phenomenon is also true for NF-κB of which activity is dependent upon activation pathway(s), posttranslational modifications, and combinatorial effects of NF-κB subunits (Cartwright *et al*, 2016). Importantly, treatment with TH5487 during OVA-induced airway hyperresponsiveness was significantly attenuated, indicative of an effect on the airway reactivity that is highly relevant for human asthma-related symptoms.

A limitation of this study is whether treatment of allergic asthma with TH5487 is feasible in humans, for which further studies are needed. Another important limitation of this study is the administration route of the drug, which is injected intraperitoneally in the mice. Due to the acute nature of allergic asthma a rapid and more directed route would be optimal, such as an inhaler. No evident changes in general murine health status were seen in this study, but more long-term monitoring of potential side effects is required before initiating clinical trials. However, previous studies on TH5487 (Visnes *et al*, 2018) and OGG1^-/-^ (Li *et al*, 2012b) mice showed no adverse health effects, with TH5487 not displaying adverse effects in AHR nor pulmonary inflammation.

Collectively, our findings display OGG1 inhibition by TH5487 as a novel, potent pharmacological approach to treat allergic asthma. TH5487 inhibition of proinflammatory genes prevents T_H_2 driven downstream activation of immune cells, resulting in reduced NF-κB activation, decreased immune cell recruitment to the lungs, lowered levels of IgE and OVA-specific IgE in plasma, reduced mucus production in the small airways, improved airway function and finally, decreased M2 macrophage populations in BALF and lung tissue. These data support further development of TH5487 and other OGG1-inhibitors as templates for novel drugs against allergic asthma.

## Acknowledgments

We would like to thank Pia Andersson for technical assistance in the lab. Graphical elements in figures were created using BioRender.com.

## Author contributions

Conceptualization: AE, LT, JB, IB, MA

Methodology: LT, JB, LP, RKVB, CD, MA, IB

Investigation: LT, JB, LP, RKVB, CD, MA, IB

Funding acquisition: AE, LT, IB, TH

Project administration: AE, LT, JB, CK

Supervision: AE, LT, JB

Writing – original draft: JB, LT

Writing – review & editing: JB, LT, RKVB, AE, IB, MA, CK, TH

## The paper explained

### Problem

Asthma is a chronic inflammatory lung disease, affecting more than 300 million people worldwide. In sensitized individuals, disease is initiated by allergenic substances triggering expression of epithelial cytokines, the primary drivers of immune responses to allergens. Primary treatment options for asthma focus on reductions in lung inflammation using inhaled beta-agonists and corticosteroids. However, a large proportion of asthma patients experience side effects, resulting in compromised treatment adherence. This necessitates the identification of novel therapeutic targets to improve patients’ disease management.

### Results

In this study, administration of TH5487 to mice with OVA-induced allergic airway inflammation significantly decreased goblet cell hyperplasia and mucus production. TH5487 treatment also decreased levels of activated NF-κB and expression of proinflammatory cytokines resulting in significantly lower recruitment of eosinophils and other immune cells to the lungs. Gene expression profiling of asthma and allergy-related proteins after TH5487 treatment revealed differences in several important regulators, including down regulation of Arg1, Mcp1 and Ccl11, and upregulation of the negative regulator of T_H_2, Bcl6. In addition, the OVA-induced airway hyperresponsiveness was significantly reduced by TH5487 treatment.

## Supplementary Materials

**Supplementary figure 1:** Whole lung scans from murine lungs stained with H&E. Scale bar = 2 mm.

**Supplementary figure 2:** Whole lung scans from murine lungs stained with PAS. Scale bar = 2 mm.

**Supplementary Figure 3: Representative images from mouse lungs showing (A) inflammatory cell influx and (B) periodic acid-Schiff (PAS) staining.** Scale bars = 100 μm and 50 μm.

**Supplementary figure 4:** Individual graphs of all cytokines measured in lung homogenate. Statistical comparison between groups was performed using a one-way ANOVA with Dunnett’s post-hoc test (\*\*\*\**P*<0.0001, \*\*\**P*<0.001, \*\**P*<0.01, \**P*<0.05).

**Supplementary figure 5:** Individual graphs of all cytokines measured in BALF. Statistical comparison between groups was performed using a one-way ANOVA with Dunnett’s post-hoc test (\*\*\*\**P*<0.0001, \*\*\**P*<0.001, \*\**P*<0.01, \**P*<0.05).

**Supplementary figure 6:** Individual graphs of all cytokines measured in plasma. Statistical comparison between groups was performed using a one-way ANOVA with Dunnett’s post-hoc test (\*\*\*\**P*<0.0001, \*\*\**P*<0.001, \*\**P*<0.01, \**P*<0.05).

**Supplementary Figure 7: TH5487-driven differences in macrophage M2 polarization in mouse lung and BALF.** (A) Western blot analysis of Arg1 and CD206 showing significant reduction of Arg1 after TH5487 treatment compared to the OVA group. (B) Cytospins from BALF samples stained with Arg1 antibodies showing an increase in Arg1 positive cell in the OVA group, which is significantly reduced by TH5487 treatment. Statistical comparisons were performed using a one-way ANOVA followed by a Dunnett’s post-hoc test (\*\*\*\**P*<0.0001, \*\*\**P*<0.001, \*\**P*<0.01, \**P*<0.05).

**Supplementary Figure 8: Representative Western blots conducted on murine lung tissue from OVA-challenged mouse lung experiments.** GAPDH was utilized as a control for all murine lung sample blots. P-RelA (ser276; MW: 67 kDa); RelA (p65; MW: 70 kDa), GAPDH (MW: 38 kDa); CD206 (MW:170 kDa); Arginase 1 (ARG 1; MW: 36 kDa).

**Supplementary Table 1:** Probes used for Real-time PCR analysis.

